# Harmonic Passive Motion Paradigm

**DOI:** 10.1101/2021.07.06.451400

**Authors:** Carlo Tiseo, Sydney Rebecca Charitos, Michael Mistry

## Abstract

How humans robustly interact with external dynamics is not yet fully understood. This work presents a hierarchical architecture of semi-autonomous controllers that can control the redundant kinematics of the limbs during dynamic interaction, even with delays comparable to the nervous system. The postural optimisation is performed via a non-linear mapping of the system kineto-static properties, and it allows independent control of the end-effector trajectories and the arms stiffness. The proposed architecture is tested in a physical simulator in the absence of gravity, presence of gravity, and with gravity plus a viscous force field. The data indicate that the architecture can generalise motor strategies to different environmental conditions. The experiments also verify the existence of a deterministic solution to the task-separation principle. The architecture is also compatible with Optimal Feedback Control and the Passive Motion Paradigm. The existence of a deterministic mapping implies that this task could be encoded in neural networks capable of generalisation of motion strategies to affine tasks.

## 1. Introduction

Animals can generate highly dexterous movements while dealing with kinematic redundancy, singularities, non-linear dynamics, uncertainties of interaction and signal delays. In contrast, all these scenarios are open problem in robotics, where available methods are not yet able to achieve animal-like robustness of interaction [1–5]. The fragility of the current model-based approaches can be related directly to the need for accurate tracking of environmental interactions to guarantee the controllers’ stability [6–8], thereby hindering the development of computational models capable of explaining the nervous system capabilities.

Impedance control has been proposed as a solution to such issues by encoding compliance into our control architectures by describing the desired behaviour in terms of equivalent dynamics [1, 9–12]. In general, identifying optimal movements and control is non-trivial and has resulted in the introduction of optimisation-based frameworks to identify desirable trade-offs. Thus, these methods require the exploitation of null-space projections and non-linear inverse optimisation that make them not robust to: singularities, model inaccuracy, information delay, and state discretisation [4–8, 13]. Bioinspired learning algorithms have also been proposed to identify desirable impedance behaviours to interact with external dynamics for specific tasks [2, 3, 14–16]. These optimised strategies, called dynamic primitives, are defined as learned dynamic attractors required to produce the desired outcome [2, 3, 17, 18]. Recently, dynamic primitives have also been proposed as an explanation to how the nervous system deals with complex dynamic interactions [17]. Primitives are classified into three categories rhythmic, discrete, and the combined behaviours. The first category includes all the movements characterised by limit cycles; thus, encoding oscillatory motions. The discrete behaviours are typical of point attractors and describe an action such as reaching [14, 17]. An example of a combined behaviour is playing drums.

Our recent work shows how adopting a hierarchical architecture of Fractal Impedance Controllers (FIC) can generate a planar human-like trajectory for upper limb reaching movements and wrist pointing trajectories [4,5]. The FIC is a well suited for this application because it allows to generate a stable, smooth behaviour without the need of using the projected inverse dynamics, which limits the controller robustness to delays and makes the system unstable when passing through singularities [4, 9, 19]. The main limitation is that most of the available control and planning algorithms rely on projected dynamics and numerical solutions to the inverse kinematics; thus, it often requires to alternative solutions to preserve such properties. This work refines and extends that method to a 7-DoF arm (Figure 1) exploiting a recently introduced geometrical postural optimisation and introducing delays into the control loop.

**Figure 1.**
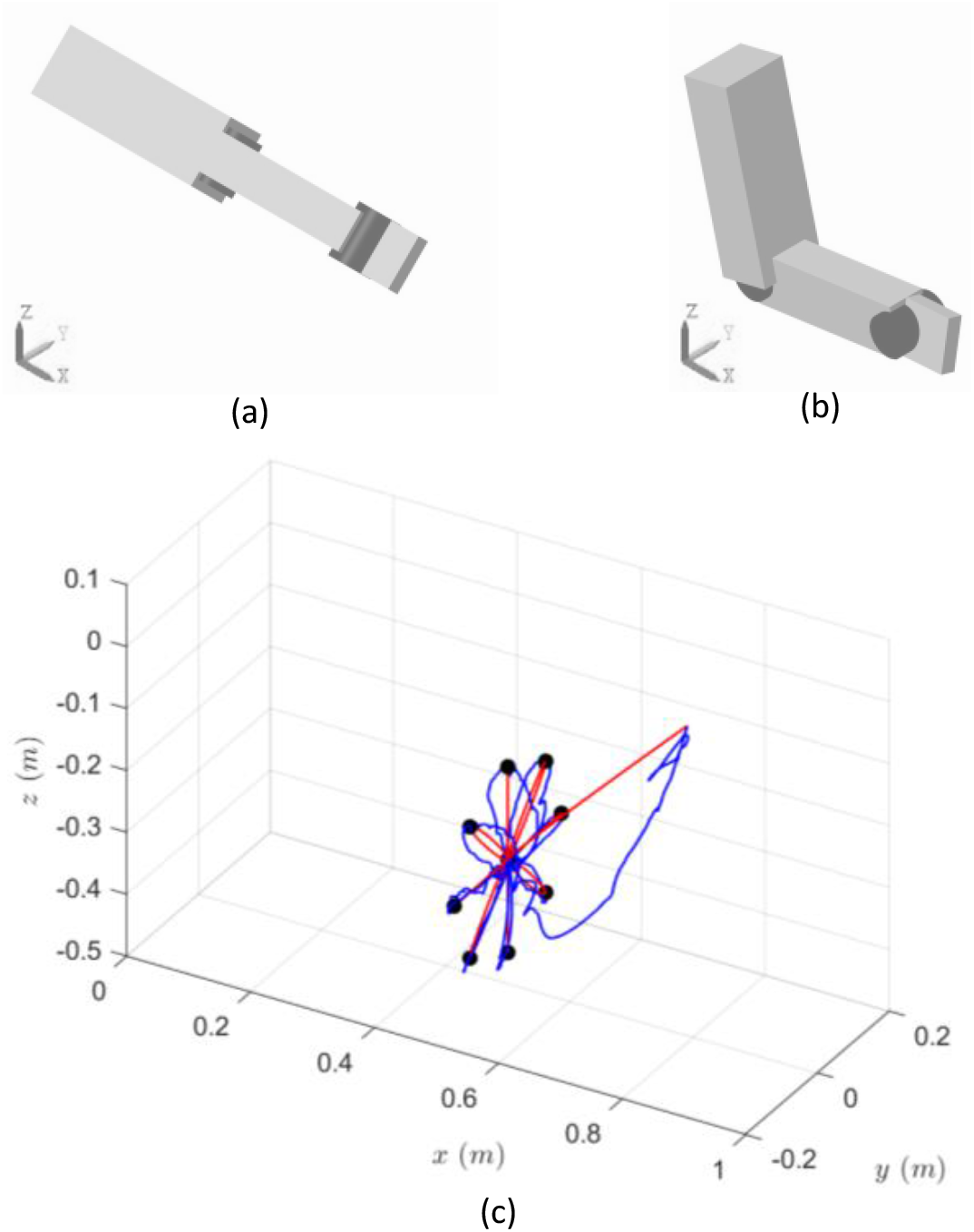
(a) Arm initial configuration in singularity. (b) Arm in the clock’s centre. (c) In red the desired trajectory, and in blue the executed trajectory in presence of gravity and a viscous force field applied at the end-effector. The black sheres are the clock’s targets.

## 2. Background on Motor Control

The Central Nervous System (CNS) can accurately control the body in extremely challenging dynamics conditions. It does so without being affected by redundancy, singularities, and by the presence of noise and delays in the sensory feedback [2, 17, 18, 20–25]. It describes Optimal Feedback Control (OFC) [26–28] and the Passive Motion Paradigm (PMP) [20,29,30]. The authors would like to remark that this section provides a general overview required for the contextualisation of the proposed control architecture. Thus, we refer the reader to the specialised literature for a detailed understanding of the neurophysiology of motor control, such as [22, 25, 31].

### 2.1. Central Nervous System

The CNS is organised with a hierarchical structure starting at the cortical level and reaching the Peripheral Nervous System (PNS) via the spinal cord. The hierarchical organisation is essential to deal with the delay involved in the neural transmission [2, 21, 22, 25, 31, 32]. For example, experimental results show that for balance tasks cortical potentials are delayed 200 − 400ms compared to en external stimulus [33], and spinal cord reflexes intervene about 100ms after a perturbation occurs [34–36].

The prefrontal cortex holds the apical position in the hierarchy of the portion of the CNS that is considered involved in motor control. It is believed to attend to the control of voluntary movements and be involved in learning new skills [25]. The basal ganglia provide the reward of performing an action by assessing the associated cost of the action. The pre-motor cortex plays a key role in solving problems associated with visual feedback and the generation of a task-space movement strategy [25, 31, 32]. The motor cortex is involved in controlling the movement dynamics, based on the proprioceptive feedback [2, 25, 32]. The parietal cortex processes the state estimation for both the proprioceptive and the visual feedback, and it is strictly interconnected with the internal models allocated in the cerebellum [2, 25, 32].

The commands issues from the motor cortex reach the Central Pattern Generators in the spinal cord, which is in charge of synchronised oscillatory movements, and they receive inputs from the motor cortex, cerebellum, and the afferent sensory information arriving from the periphery [2, 25, 37, 38]. The spinal cord is also the loci of the spinal cord reflexes that are the first intervention strategy in the spinal cord. These reflexes provide the first centralised response to proprioceptive feedback [2,21,36,37]. It is worth noting that considering that the delays of these reflexes are in the order of 100 ms, the modulation of the muscle-skeletal system dynamics is essential to movement stability by filtering high-frequency perturbations.

### 2.2. Optimal Feedback Control & Separation Principle

The OFC exploits optimal control theory to explain how the nervous systems optimise human movements, stabilise the task, and control the redundant degrees of freedoms [26, 27, 31]. The OFC describes the nervous systems as a model-based continuous feedback controller, which exploits multiple cost functions for different tasks [28, 39]. The framework was successfully used to model the separation principle by classifying these controllers for dynamic and static tasks [20, 28, 30, 39]. Dynamic controllers deal with the interaction with the dynamics forces (inertial and velocity-dependent forces), and the static controllers compensate for the static forces (elastic fields and gravity) [28]. The model has been proven capable of mimicking human behaviour, capturing how motor control adjusts to sensory feedback during task execution [28, 39]. However, the nervous system deals with the non-holonomic manifolds generated from their joint-space formulation, which is not compatible with the holonomic characteristics experimentally observed in humans and described by Donders’Law [20, 30, 31].

### 2.3. Equilibrium Point Hypothesis & Passive Motion Paradigm

A well-known theory for movement generation is the Equilibrium Point Hypothesis (EPH) which describes the generation of reaching movements as a gradual shift of the equilibrium point [2, 20, 29, 30, 40]. Further evidence also shows that the brain process separately deals with postural control (i.e., static forces) and motion dynamics, known as the separation principle. The PMP is a control framework that utilises this principle [2, 4, 5, 20, 30]. The *λ*_0_-PMP is a recently proposed extension of this framework. It relies on a viscous force field to produce a constraint that limits the projections of some of the joint torques in the task space while optimising posture in the manipulator null-space [20,30]. Other studies show that a similar result can be obtained by superimposing multiple task-space conservative non-linear impedance controllers to generate virtual mechanical constraints along the kinematic chain. This approach relies on a Fractal Impedance Controller (FIC) to define a non-linear impedance controller around the desired state [4, 5, 41]. Differently from the *λ*_0_-PMP this latest approach has the advantage that it does not rely on identifying a damping component and matrix inversions, which is taken over by the introduction of a non-linear spring [5]. Therefore, the controller energy is a path independent potential field (i.e., holonomic); thus, respecting Donders’s Law [5, 20, 30].

## 3. Harmonic Passive Motion Paradigm

The Harmonic Passive Motion Paradigm (H-PMP) is a controller architecture describing communication from the motor cortex to its related muscle groups. The H-PMP can integrate haptic control, but this functionality is omitted for simplicity in this work. The H-PMP architecture is schematised in Figure 2.

**Figure 2.**
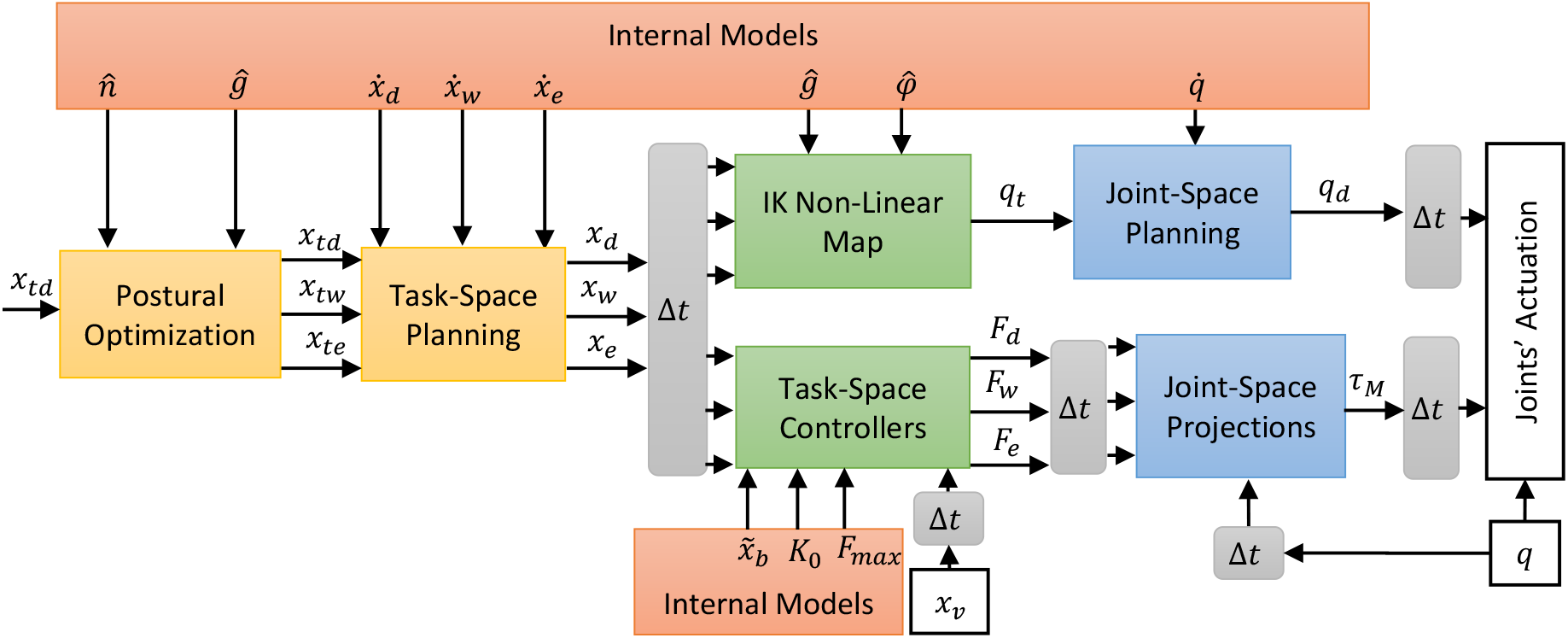
The H-PMP architecture starts from the postural optimisation of the limb. It takes into input the end-effector target for the hand *x*_td_, and an internal model of the task in terms of motion plane 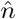 and grasp direction *ĝ*. The postural optimisation generates targets for the wrist (*x*_w_) and the elbow (*x*_e_). These three targets are the input of the task-space planners, which based on the desired velocities associated with the task’s internal model (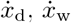 and 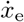 generate the limbs’ trajectories. The limbs’ trajectories are delayed on an interval Δ*t* and, subsequently, passed as input to both the IK and the task-space controllers. The IK is solved using a non-linear map which depends from 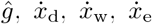, and the unit vector for wrist pronosupination 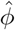, and its output is passed to the joints’ trajectory planners to derive the desired joints’ trajectories (*q*_d_). The three task space controllers input are the desired end-effector trajectories, their delayed visual feedback (*x*_v_), and the models’ parameters of the non-linear impedance (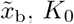, and *F*_max_). The output forces are passed to derive the desired joints’ torques (*τ*_M_), which requires the knowledge of the joints’ configuration *q*. Lastly, the *q*_d_ and *τ*_M_ are the input of the joints’ actuation modelling the coordinated actions of the muscle groups.

### 3.1. Cartesian-Space Harmonic Trajectory Planning

Cartesian trajectory planning is traditionally associated to a Cost-To-Go of the basal ganglia, usually implemented via optimisation to generate minimum jerk trajectories. We have recently proposed a planner which employs the FIC to generate harmonic trajectories that have been proven to capable of generating human-like kinematics on planar arm kinematics in [4,5,13,42]. Harmonic trajectories are preferable because they require a lower peak power compared to minimum jerk trajectories for generating the same average velocity [4, 5]. An independent planner for each of the task-space DoF is implemented as an Model Predictive Controller (MPC) per unit of mass. The desired trajectory is computed via the integration of the desired acceleration 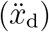 output from the controller.

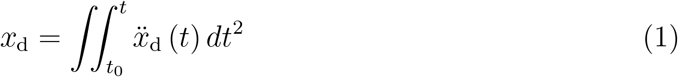

where *t*_0_ is the initial time, and *t* is the current time. The acceleration is derived from the following equation:

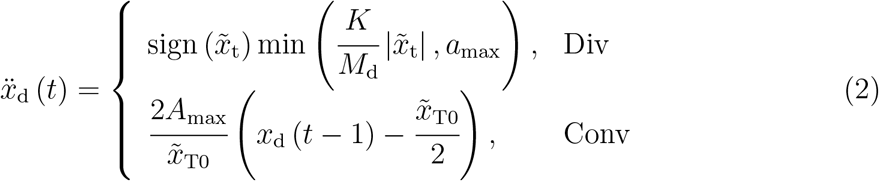

Where 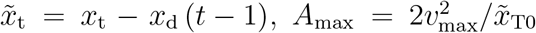 is the acceleration computed at the maximum displacement 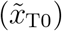 reached in the previous divergence phase, and *v*_max_ = 1.596*v*_d_ is the maximum velocity of the harmonic trajectory generating the desired velocity *v*_d_.

### 3.2. Arm Postural Optimisation

The arm postural optimisation and IK exploit a recently introduced geometrical solution, described by one of the functions of the motor cortex. We have chosen this approach because it generates a deterministic non-linear mapping robust to singularities between the task-space and the joint-space exploiting the kineto-static duality [43], and could be performed by the nervous system relying on an internal model present in the cerebellum. The definition of the optimal posture can be identified by aligning the arm plane of motion with the task dynamics, leading to manipulability properties equivalent to a planar 3-link arm [43].

Let us define the following unit vectors requires by the formulations of the postural optimisation and inverse kinematics:

- 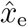 is the direction of the arm link
- 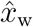 is the direction of the wrist vector
- 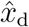 is the direction of end-effector vector
- *ĝ* is the pointing direction
- 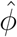 is the direction controlling the hand-pronosupination
- *l*_a_ is the arm length
- *l*_fa_ is the forearm length
- *l*_h_ is the hand length
- 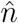 as the unit vector orthogonal to the task’s plane

The postural optimisation equations for a left-arm can now be written as follows:

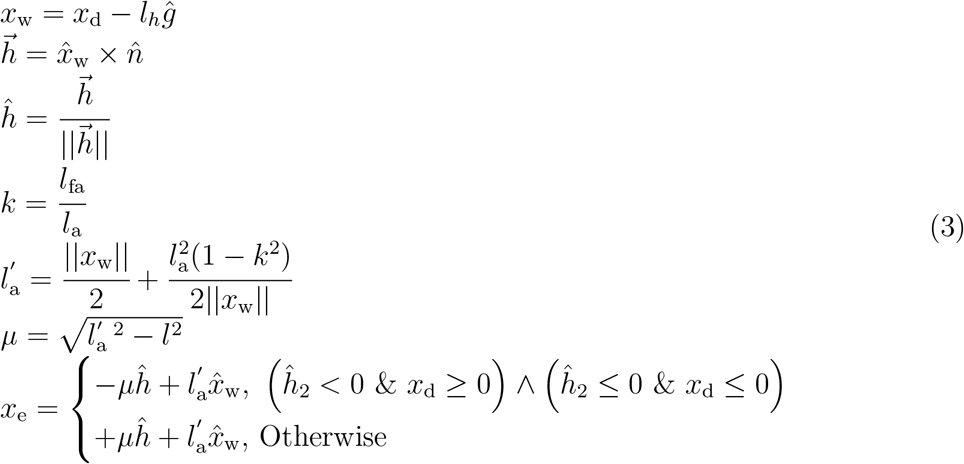

The equation for the right arm can be easily obtained by inverting the signs of *µĥ*. The joint angle can be derived from the following equations:

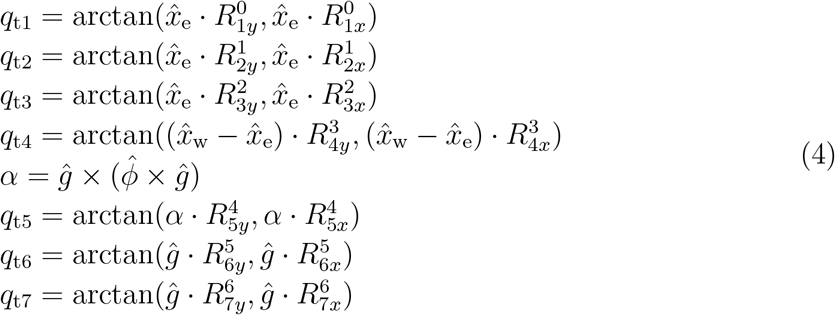

where *q*_t*i*_ is the *i*^*th*^ joint angle of the target, *R*_*i*_ is the rotation matrix of the base from of the *i*^*th*^ link, and 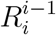 is the base frame of the *i*^*th*^ joint.

### 3.3. Joint-Space Harmonic Trajectory Planning

Joint space planners are based on a variation of the model used for the Task-Space planning in subsection 3.1. The maximum angular acceleration and velocity are descriptive of the maximum mechanical characteristics of each joint.

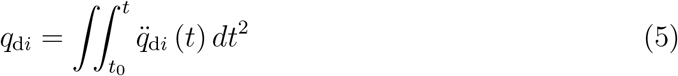

The acceleration is derived from the following equation:

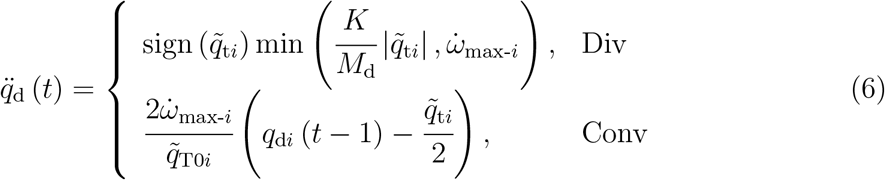

where 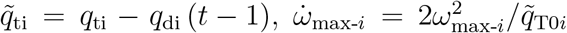 is the maximum acceleration that can be computed known the maximum angular velocity (*ω*_max-*i*_) and the maximum displacement reached in the previous divergence phase 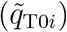.

### 3.4. Task-Space Interaction Control

The Task-Space interaction is a variation superimposition of multiple Fractal Impedance controllers based on the methods tested in [4, 41]. Task-space controllers are assigned to control the pronosupination and each of the 3-links composing the arm. They will prescribe the effort along the kinematic chain determining the response to external dynamics and mechanical losses. The projection of such information in joint space will be explained in the next section.

The FIC effort (i.e, force or torque) is computed according to the following equation:

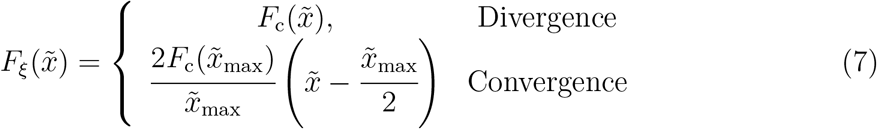

where *F*_c_ is a generic continuous and upper-bounded force profile, 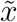 is the pose error, 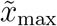 is maximum pose error reached during the previous divergence phase [4, 11, 42]. The chosen force profile is single sigmoidal force saturation enclosing a region of linear impedance around the desired pose, based on the formulation proposed in [4, 42].

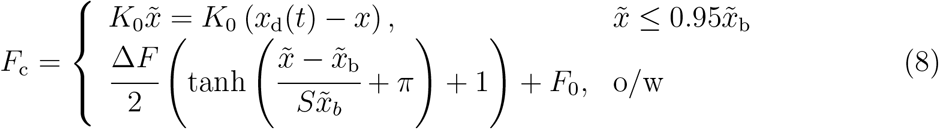

where *K*_0_ is the constant stiffness, 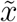 is the end-effector pose error, Δ*F* = *F*_max_ − *F*_0_, 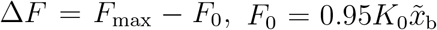, and *S* = 0.008 controls the saturation speed to be completed between the last 5 % of 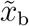.

### 3.5. Joint-Space Projections

Joint coordination is believed to be addressed via the central pattern generators of the spinal cord, which is also responsible for the control of the reflexes due to its proximity to the periphery. This role is modelled in the proposed method as a geometric projection of the desired task-space forces determined by the task-space controllers shown in Figure 2. Additionally, We have allocated this logic for the generation of the torque commands a low level, to reduce the delay in the transmission of postural information used to compute the Jacobians. It is worth noting that although we have omitted the reflex in this work for simplicity, they could be implemented using the haptic algorithm presented in [42]. The desired forces *F*_*ξ*_ from Equation 7 are computed for the elbow *F*_e_, the wrist *F*_w_, and the end-effector *F*_d_ before projecting them in the joint space using the respective geometric Jacobians (*J*_e_, *J*_w_ and *J*_d_).

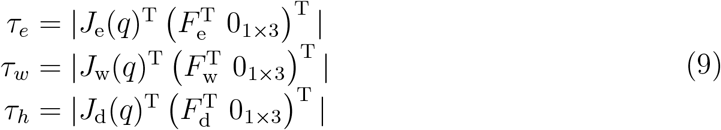

Meanwhile the pronosupination control directly generates a torque *τ*_PS_ acting only on the fifth joint controlling the pronosupination. The four torques vectors are then combined to control the maximum torque of the joints’ actuation presented in the following section.

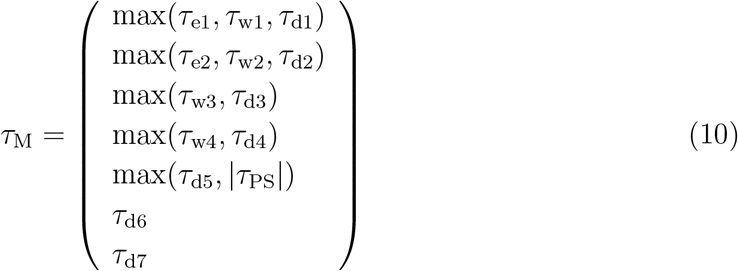

where *τ*_ei_, *τ*_wi_, *τ*_di_ are the *i*^*th*^ components of the respective vectors. This strategy guarantees the maximum rigidity of the arm based on the output of the 4 independent task-space controllers (3 links plus wrist pronosupination).

### 3.6. Joints’ Controller

The proposed control architecture describes the overall joint actuation resulting from the muscle activity in each joint. These controllers coupled with the torque projections described in subsection 3.5 models the muscle groups, which are coordinated muscles that are synchronously activated to generate movement strategies and they can influence multiple joints [44]. Therefore, each of the joints are controlled using a fractal impedance controller generating the following joint command:

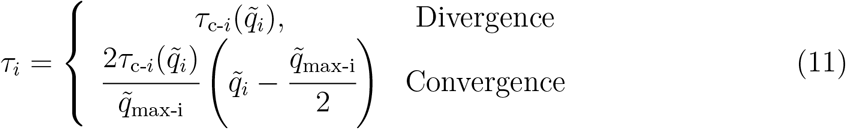

where *τ*_c-*i*_ is a continuous upper-bounded torque profile, 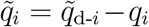 is the error between the current and the desired angle, 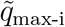 is the maximum angular error reached during the previous divergence phase. The torque profile (*τ*_c-*i*_) used in this paper is:

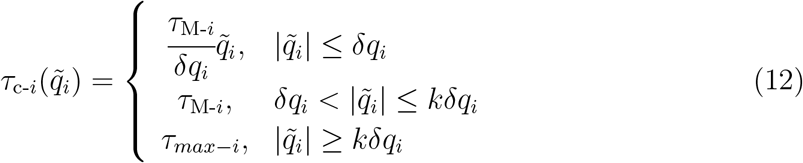

where *τ*_*max*−*i*_ is the maximum actuation torque, *δq*_*i*_ is angular accuracy, and *k* a scaling factor. The chosen formulation allows to model the muscular activities as a change of the equilibrium position of a potential field generated by the agonist and antagonist muscles. This choice is based on theoretical and experimental data showing that the antagonist muscles group are responsible for the modulation of the joints’ impedance [4, 45].

### 3.7. Comparison of H-PMP with other Theories

The things that OFC and the PMP approach to motor control have in common are that they are trying to describe the optimisation and stability of human movement by looking at the system’s energy. However, they are choosing different representations for the system energy, which affects both the stability of the problem and its numerical complexity.

The OFC relies on the more general formulation Optimal Control Theory. Therefore, the representation of the energy is detached the physical meaning of the energy function (i.e., cost function) and does not necessarily consider the simplification introduced by intrinsic characteristics of the physical world [26–28]. For example, does not a priory consider that Newtonian physics is limited to Euclidean geometry; therefore, if the system dynamics can be described in Newtonian physics, the energy can be regarded as *a priori* bounded to that geometry. On the contrary, the Optimal Control Theory is a general framework that can rely on the more general formulation for the metric space. However, this formulation requires the introduction of constraints to bind the solution to the physical property of the world. The non-linear dynamic constraints of the task-space projection in the generalise coordinate (i.e., joint space) requires introducing the projected dynamics into the problem formulation [6–8]. Therefore, resulting in a numerical problem that is susceptible to singularity, and the identified solution are path-dependent and cannot be generalised.

The PMP methods use a subset of Optimal Control Theory bounding the representation to a mechanical impedance, which implies that the problem formulation is bounded to the Riemannian Geometry of the system Lagrangian (i.e., energy) in generalised coordinate (i.e., joint-space) [20, 29]. The main problem now is to identify the equivalent dynamics of the system for any given task, which is mostly resolved using projected dynamics leading to the same limitations of the OFC. For example, [20] relies on a non-linear inverse optimisation to identify the *λ*_0_ term of the equation that is used to introduce the task-separation principle.

The main benefit of the H-PMP is that it does not require any explicit formulation of the system dynamics to guarantee stability, and it relies on geometric projections based on the kineto-static duality for going back and forward from between the joint and task space. The generated algorithm is computationally efficient because it does not rely on numerical optimisation and projected dynamics. This is possible because the dynamics constraints of the system’s physical properties are embedded in the controller, and the trajectory exploits the physical space fabric to determine the optimal trajectory for energy transport (i.e., harmonic trajectory). It is worth reminding that physical systems are Riemmanian manifolds, which implies that by definition, the optimal way to transfer energy is using harmonic oscillations, and any trajectory can be described as a superimposition of harmonics [46]. However, using the proposed approach requires an architecture that relies on only the projection of efforts (force and torques) from the task to the joint space and, consequently, only the projection of kinematics from the joint to the task space. Additionally, it also requires the identifications of geometrical solutions that are designed *ad hoc* on the system mechanism [42, 43]. Essential to solving this problem is the FIC because it provides an algorithmic representation of a stable, conservative non-linear oscillator that can be online tuned to oscillate with a harmonic trajectory at the desired frequency and, consequently, control the movement velocity. This characteristics are also preserved across non-linear projections [5, 42], and in presence of delays [9, 10].

To summarise, the proposed architecture in Figure 2 can be seen as an algorithmic representation of a system composed of coordinated adjustable oscillators efficient to run on silicon, which moves between equilibrium states using harmonic trajectories. The hypothesis is that the motor commands arise thanks to a similar architecture where the synchronisation occurs between neural circuits that behave as non-linear adjustable oscillators rather than between Fractal Impedance Controllers.

## 4. Validation Method

The H-PMP has been validated on two arm simulators in Simulink (Mathworks, USA) developed using the Simscape library for physics simulations, using the ode-45 solver with a maximum time step of 1 s. The first simulator evaluates the proposed method without accounting for the information delay. The second simulator evaluates the H-PMP performance in the presence of delays comparable with the nervous system without using a state estimator.

### 4.1. Arm Model

The arm is modelled as a 7-dof manipulator with a spherical-revolute-spherical joint configuration. The shoulder joint has a ZYX configuration; meanwhile, the wrist joint has an XYZ configuration. The lengths of the arm, forearm and hand are *l*_a_ = 0.37 m, *l*_a_ = 0.32m and *l*_h_ = 0.1m, respectively. Their masses are *M*_a_ = 4.1 kg, *M*_fa_ = 2.4 kg, and *M*_h_ = 1 kg. Each joint has a friction coefficient of *K*_*µ*_ = 0.5 Nms/ deg.

#### 4.1.1. Planner & Controller Parameters

The task-space planners have been set with a reference velocity *v*_d_ = 0.5 ms^−1^ with a minimum *A*_max_ = 0.1 m/s^2^, and *K/M*_d_ = 4000 s^−2^ The joint-space planners have been set with a reference velocity *ω*_d_ = 0.1*π* rad s^−1^, and *K/M*_d_ = 4000 s^−2^. The pronosupination unit vector 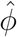 is fixed at (0 0 1)^*T*^ for all experiments. Similarly the pointing unit vector *ĝ* is also been kept constant at (1 0 0)^*T*^ for all the experiment.

The parameters of the hand’s task-space controller are *K*_0_ = 10 000 Nm^−1^, 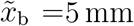, and *F*_max_ = 200 N. The parameters of the wrist’s task-space controller are *K*_0_ = 6000 Nm^−1^, 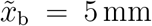, and *F*_max_ = 200 N. The parameters of the elbow’s task-space controller are *K*_0_ = 1000 Nm^−1^, 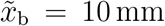, and *F* max = 100 N. The torsional reference pronosupination torque *τ*_*PS*_ has been kept constant at 15 Nm, being 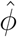 constant. The parameters of the joints’ controllers are *k* = 5, *δq*_*i*_ = 0.0175 rad ∀i ∈ [1, 7], and *τ*_*max*_ = (300 300 150 150 50 50 50)^*T*^ Nm.

#### 4.1.2. Information Delays

The Δ*t* used between the task-space planning and the IK against the task-space controller in Figure 2 is 100 ms. The delay introduced between the joint-space planning and the joints’ actuation is also 100 ms. The delay between the task-space controllers and the joint-space projections is 40 ms. Consequently, the delay between the latter component and the joints’ actuation is 60 ms. The delay of the visual feedback (*x*_v_ in Figure 2) has been set to 200 ms, and the delay for the proprioceptive feedback to the joint-space projections is 40 ms.

### 4.2. Clock Experiment

The clock experiment experiment is a common tool used for studying human movements and the motor control framework [20, 28, 30, 47]. The centre of the clock is in (0.500 0 − 0.300)m and has a radius of 15 cm. Thus, the coordinates of the 8 targets are: (0.500 0.106 − 0.406)m, (0.500 0 − 0.450)m, (0.500 − 0.106 − 0.406)m, (0.500 − 0.150 − 0.300)m, (0.500 − 0.106 − 0.194)m, (0.500 0 − 0.150)m, (0.500 0.106 − 0.194)m, and (0.500 0.150 − 0.300)m.

The experiment has been performed three times for each simulator to evaluate the effect on external force fields on the arm stability, and the tracking performances. All the simulations started from a fully extended posture aligned with the x-axis. The three conditions are: without gravity, with gravity, and with gravity plus viscous force field applied to the hand’s end-effect.

The formulation of the viscous force field is:

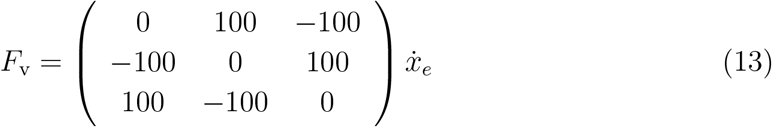

### 4.3. Data Analysis

The Mean Absolute Error (MAE) and the Root Mean Square Error (RMSE) are computed for the trajectories of the task-space and joint-space controllers. The MAE is used to evaluate the average tracking accuracy. Meanwhile, the RMSE quantifies the impact of sporadic large errors (i.e., outliers) on the tracking performance.

The mean and the standard deviation of the ratio between the average and the peak velocity 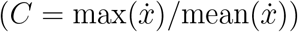 of the task-space trajectories have been computed and compared with the human range of *C*_H_ = 1.805±0.153 [4,40,48] to validate the similarity with human movements.

## 5. Results

The planned and the executed trajectories in Figure 3 show that the arm always reaches, but there is a reduction of the tracking accuracy with the perturbation increase. The MAE and the RMSE are reported in Table 1. The RMSE is higher, suggesting the sporadic occurrence of large errors during the trajectory tracking. The two tracking errors increase when introducing the delays, which is expected due to the increased time-shift between the issuing of the command and its execution. The data also show that the accuracy at the elbow end-effector is constrained to 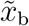, but it degenerates beyond 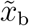 for both the wrist and the hand controllers. Thus, suggesting that it might be related to the accumulation of the joints’ position errors and can be compensated by introducing a state estimator. The MSA and the RMSA in the joint-space (Table 2) show that the errors of the first five joints are always constrained within the accuracy set in the controllers. Meanwhile, the sixth and the seventh joints are consistently beyond suggesting a need for an increase in the system stiffness to achieve the desired accuracy. The joints’ errors also confirm that the task-space accuracy can be improved by introducing a state estimator to compensate for the systematic error.

**Figure 3.**
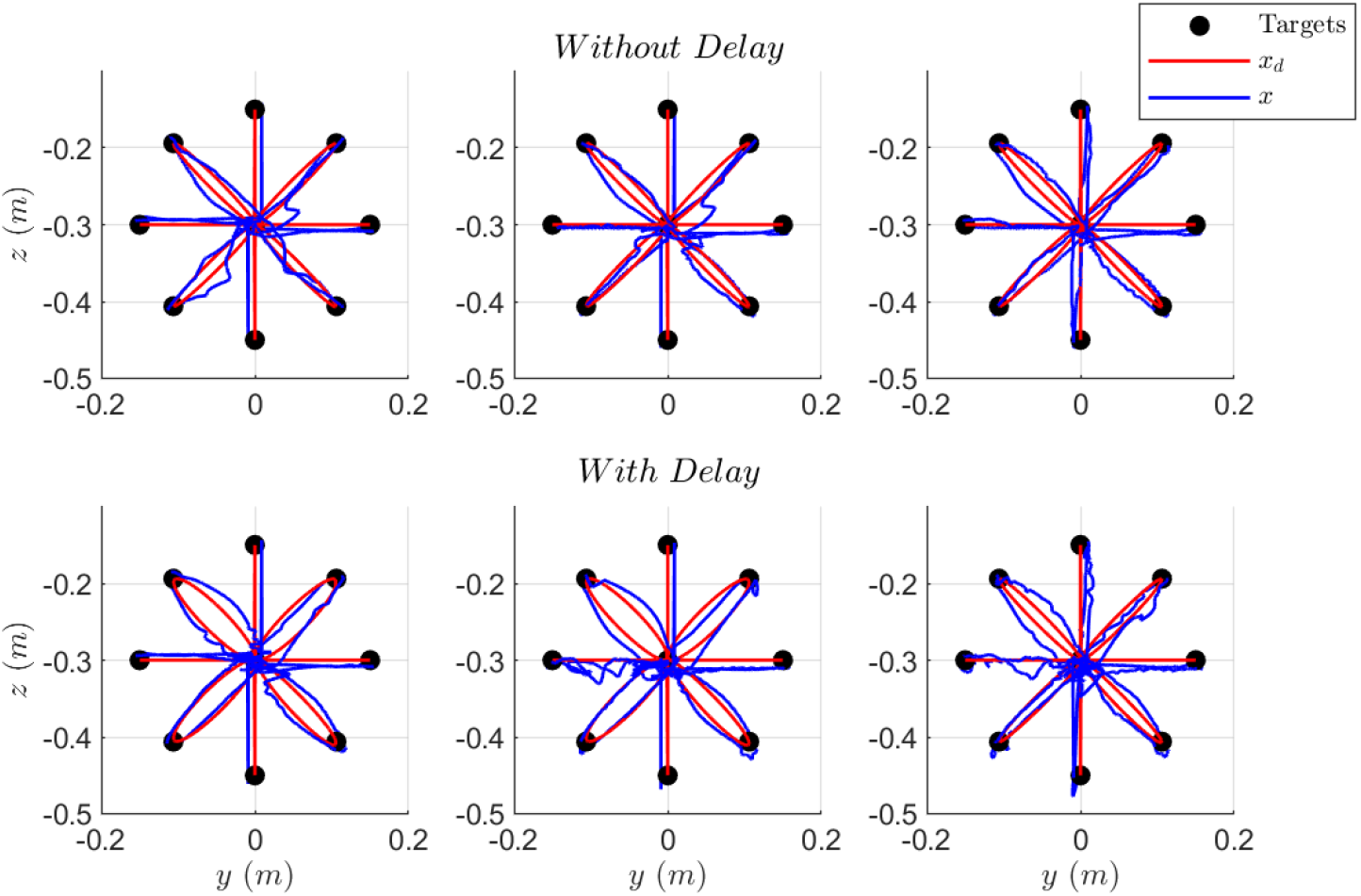
Reaching trajectories in the three considered conditions for the H-PMP with and without delay. The trajectories in the absence of gravity are on the left. The data recorded for the gravity scenario are in the centre. The trajectories for the scenario of gravity plus viscous force field are on the right. We observe a general reduction of accuracy moving from right to left.

**Table 1.** The MEA and RMSE for the task-space controllers.

| | MAE($\tilde{x}_e$) [mm] | MAE($\tilde{x}_w$) [mm] | MAE($\tilde{x}_d$) [mm] |
| --- | --- | --- | --- |
| No Gravity | (0.3 0.2 1.1)<sup>T</sup> | (6.8 2.3 3.4)<sup>T</sup> | (6.8 2.3 3.4)<sup>T</sup> |
| Gravity | (1.6 0.4 1.5)<sup>T</sup> | (6.5 1.3 4.5)<sup>T</sup> | (6.5 1.6 4.5)<sup>T</sup> |
| Viscous Field | (1.7 0.6 1.5)<sup>T</sup> | (6.5 1.6 4.7)<sup>T</sup> | (6.5 1.6 4.7)<sup>T</sup> |
| No Gravity & Delay | (1.5 3.6 1.8)<sup>T</sup> | (7.5 6.0 6.0)<sup>T</sup> | (7.5 6.0 6.0)<sup>T</sup> |
| Gravity & Delay | (2.5 3.7 2.1)<sup>T</sup> | (7.5 6.1 7.5)<sup>T</sup> | (7.5 6.1 7.5)<sup>T</sup> |
| Viscous Field & Delay | (2.7 3.8 2.1)<sup>T</sup> | (6.8 6.4 8.0)<sup>T</sup> | (6.8 6.4 8.0)<sup>T</sup> |
| | RMSE($\tilde{x}_e$) [mm] | RMSE($\tilde{x}_w$) [mm] | RMSE($\tilde{x}_d$) [mm] |
| No Gravity | (0.8 0.6 3.0)<sup>T</sup> | (10.6 5.6 5.3)<sup>T</sup> | (10.5 8.9 7.8)<sup>T</sup> |
| Gravity | (2.1 0.7 3.1)<sup>T</sup> | (10.5 2.8 6.0)<sup>T</sup> | (10.5 8.0 9.6)<sup>T</sup> |
| Viscous Field | (2.2 1.3 3.2)<sup>T</sup> | (10.4 2.4 6.3)<sup>T</sup> | (10.4 7.2 9.4)<sup>T</sup> |
| No Gravity & Delay | (3.6 6.6 3.6)<sup>T</sup> | (11.5 8.6 8.3)<sup>T</sup> | (11.5 11.9 8.4)<sup>T</sup> |
| Gravity & Delay | (4.1 6.6 3.8)<sup>T</sup> | (11.4 8.8 9.5)<sup>T</sup> | (11.4 12.0 10.9)<sup>T</sup> |
| Viscous Field & Delay | (4.5 6.6 3.8)<sup>T</sup> | (10.4 8.9 10.2)<sup>T</sup> | (10.4 12.3 11.4)<sup>T</sup> |

**Table 2.** The MSA and RMSE for joint-space controllers.

| | MAE($\tilde{q}$) [mrad] |
| --- | --- |
| No Gravity | (2.2 0.0 8.3 6.3 2.1 69.4 76.4)<sup>T</sup> |
| Gravity | (1.3 4.9 3.9 5.8 2.1 69.3 76.5)<sup>T</sup> |
| Viscous Field | (3.8 5.3 4.9 6.6 2.1 69.7 72.0)<sup>T</sup> |
| No Gravity & Delay | (1.6 0.5 3.7 1.8 2.2 69.4 76.5)<sup>T</sup> |
| Gravity & Delay | (1.4 4.7 3.7 4.9 2.2 70.5 76.6)<sup>T</sup> |
| Viscous Field & Delay | (2.5 5.1 3.2 5.0 2.2 69.1 71.9)<sup>T</sup> |
| | RMSE($\tilde{q}$) [mrad] |
| No Gravity | (7.0 0.7 20.7 11.8 3.4 72.7 79.6)<sup>T</sup> |
| Gravity | (2.9 6.1 9.9 9.7 3.4 74.7 79.6)<sup>T</sup> |
| Viscous Field | (10.4 6.7 12.3 10.6 3.4 73.9 76.2)<sup>T</sup> |
| No Gravity & Delay | (4.4 1.0 9.3 3.6 3.5 72.7 79.7)<sup>T</sup> |
| Gravity & Delay | (3.6 7.1 11.2 9.0 3.5 75.1 79.7)<sup>T</sup> |
| Viscous Field & Delay | (5.5 8.2 6.8 9.0 3.5 73.5 76.0)<sup>T</sup> |

The distribution of the *C* values for the planned trajectory (Figure 4) are consistent with the value of the harmonic trajectories (*C*_HT_ = 1.596), and the distribution have little variance which does not overlap with the interval measured for human movements *C*_H_ = 1.805 ± 0.153 [40]. Figure 5 presents the data for the executed trajectories showing a substantial overlap with *C*_H_. The One-Way ANOVA has shown no significant difference in the C values recorded in the six experimental conditions. The test *p*-value is 0.9442. The data considered as belonging to a single population are compared with the human data in Figure 6, which confirms the overlap. The data in Figure 5 and Figure 6 show that the data from our simulation show a higher variability compared to both the human data and our preliminary results obtained for a 3-link planar arm [4]. It indicates that this variability could be related to sudden velocity peaks due to perturbation or not optimal coordination between the different architectural components. This analysis is confirmed by the data on the end-effector tangential velocity in Figure 7, showing that the executed trajectories have an expected trend of a 10 to 15% increase in peak velocity compared to the planned trajectories. However, the data also show that there are episodically peaks that are substantially higher, which supports the hypothesis mentioned above that they are related to local conditions due to perturbations and lack of coordination.

**Figure 4.**
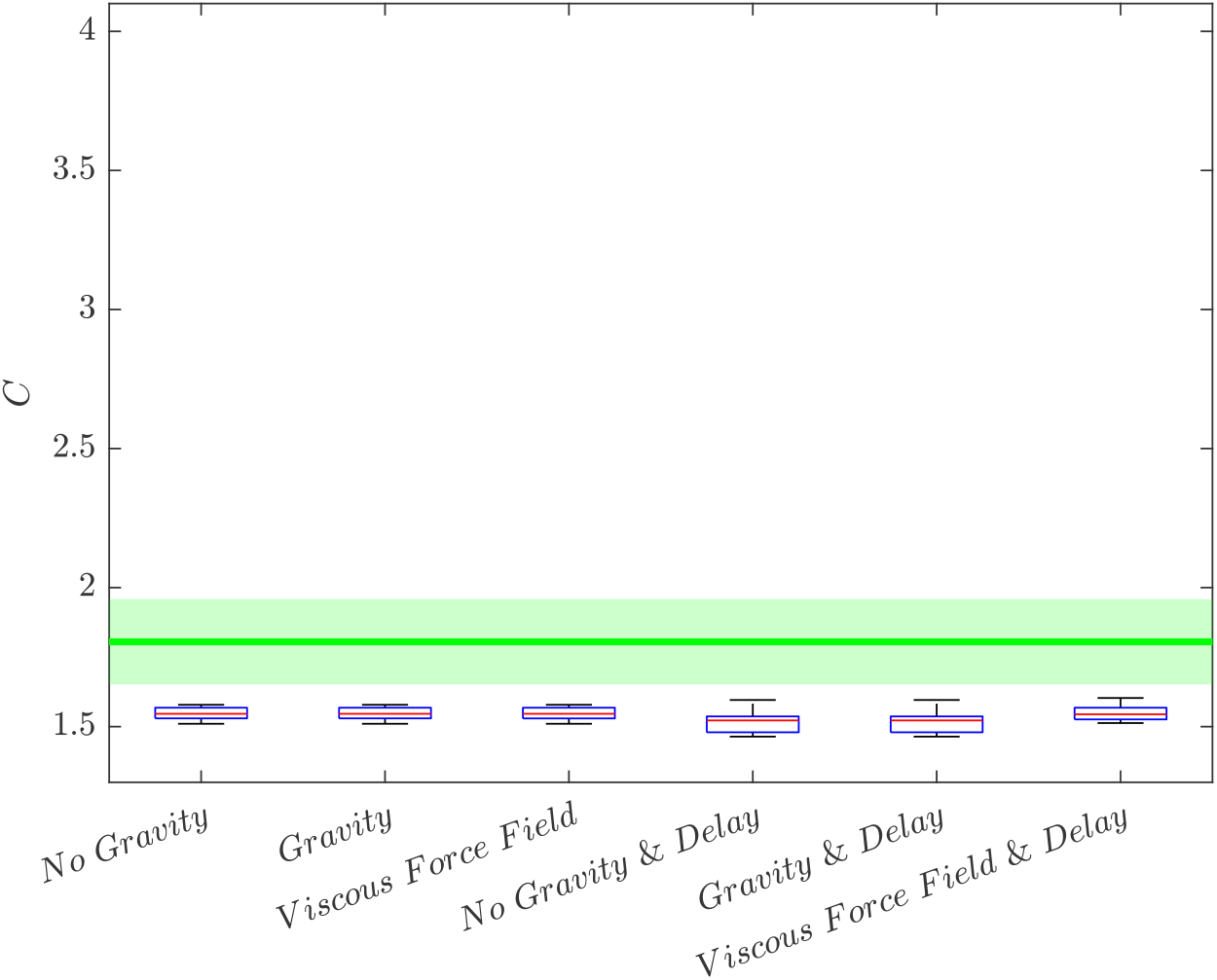
The *C* value of the planned trajectories is consistent with harmonic trajectories (*C*_HT_ = 1.596) [4], which is significantly lower compared to the human values of *C*_H_ = 1.805 ± 0.153 [40, 48], reported in green.

**Figure 5.**
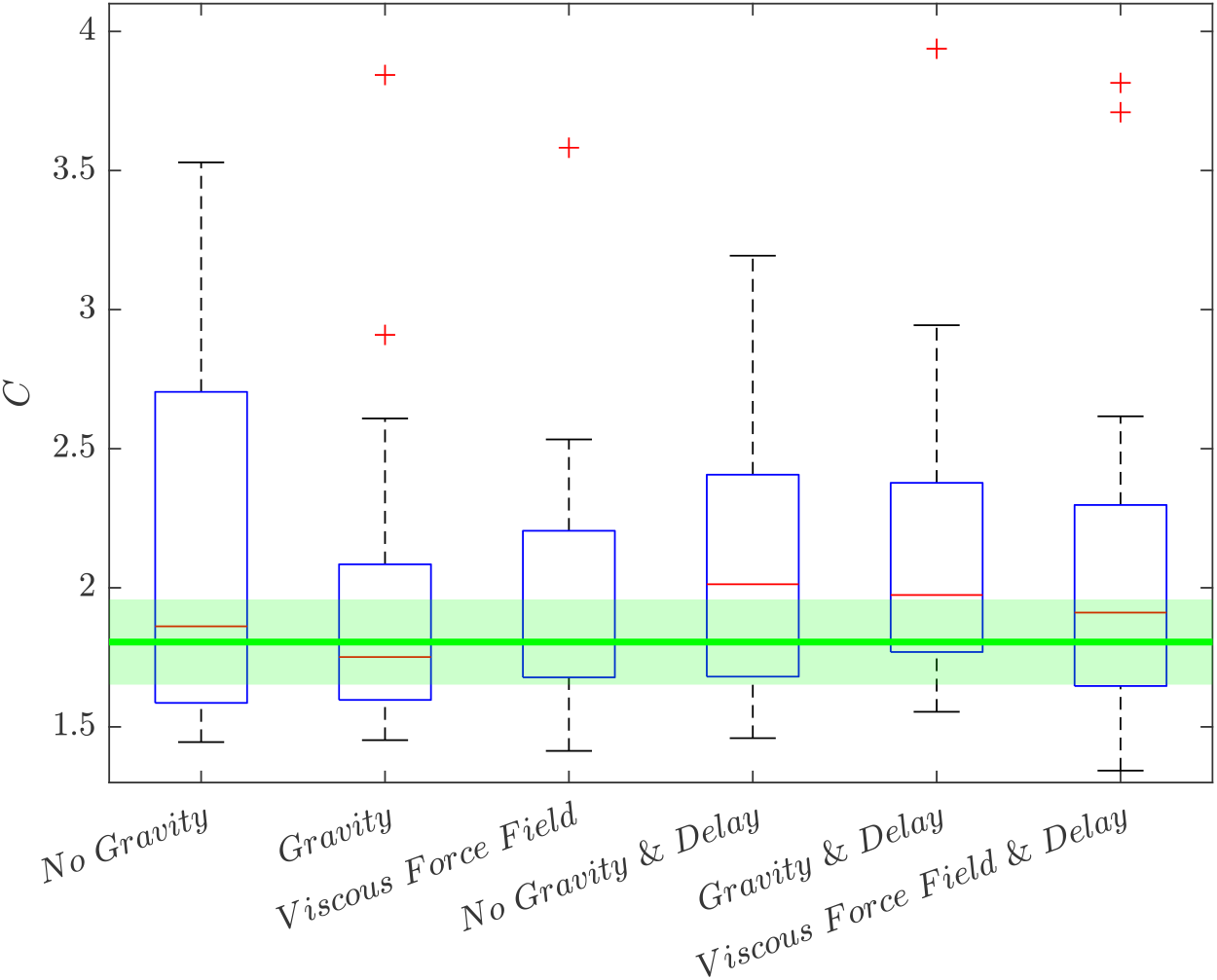
The *C* value of the executed trajectories increases compared to the values of the planner trajectories. The data show high variability of the values probably connected to velocity peaks due to local perturbation, considering the tracking errors reported in Table 1. The human values of *C*_H_ = 1.805 ± 0.153, [40], are reported in green, and their mean value is contained within the standard deviation in all the experimental conditions.

**Figure 6.**
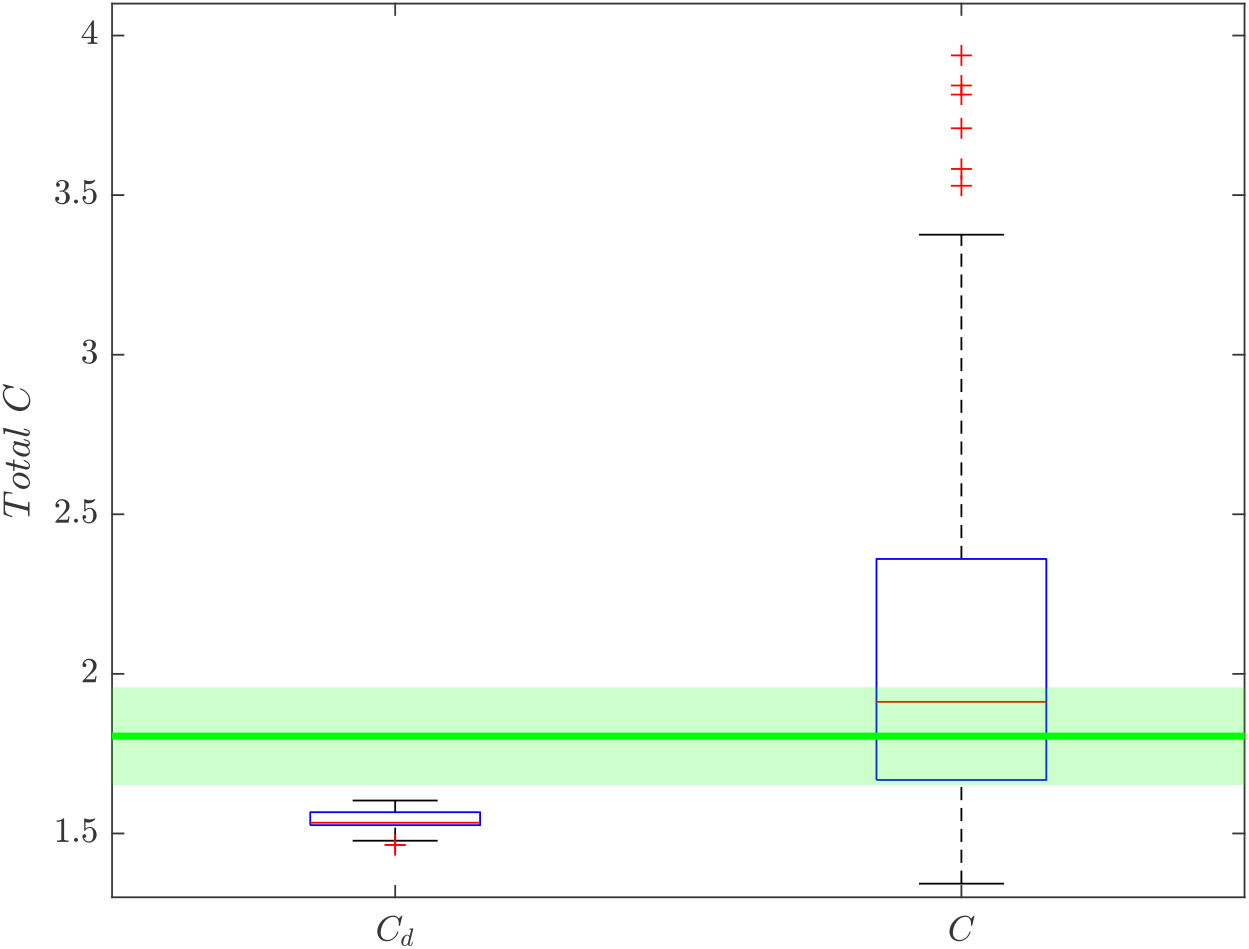
*C*_d_ are the values from the planner, *C* indicates the values for the executed trajectories. The One-Way ANOVA confirmed that the C values of the measured trajectories from the different experimental condition are from the same population (*p* = 0.9442). The human values of *C*_H_ = 1.805 ± 0.153, [40], are reported in green. Confirming a substantial overlap of the trajectories distributions.

**Figure 7.**
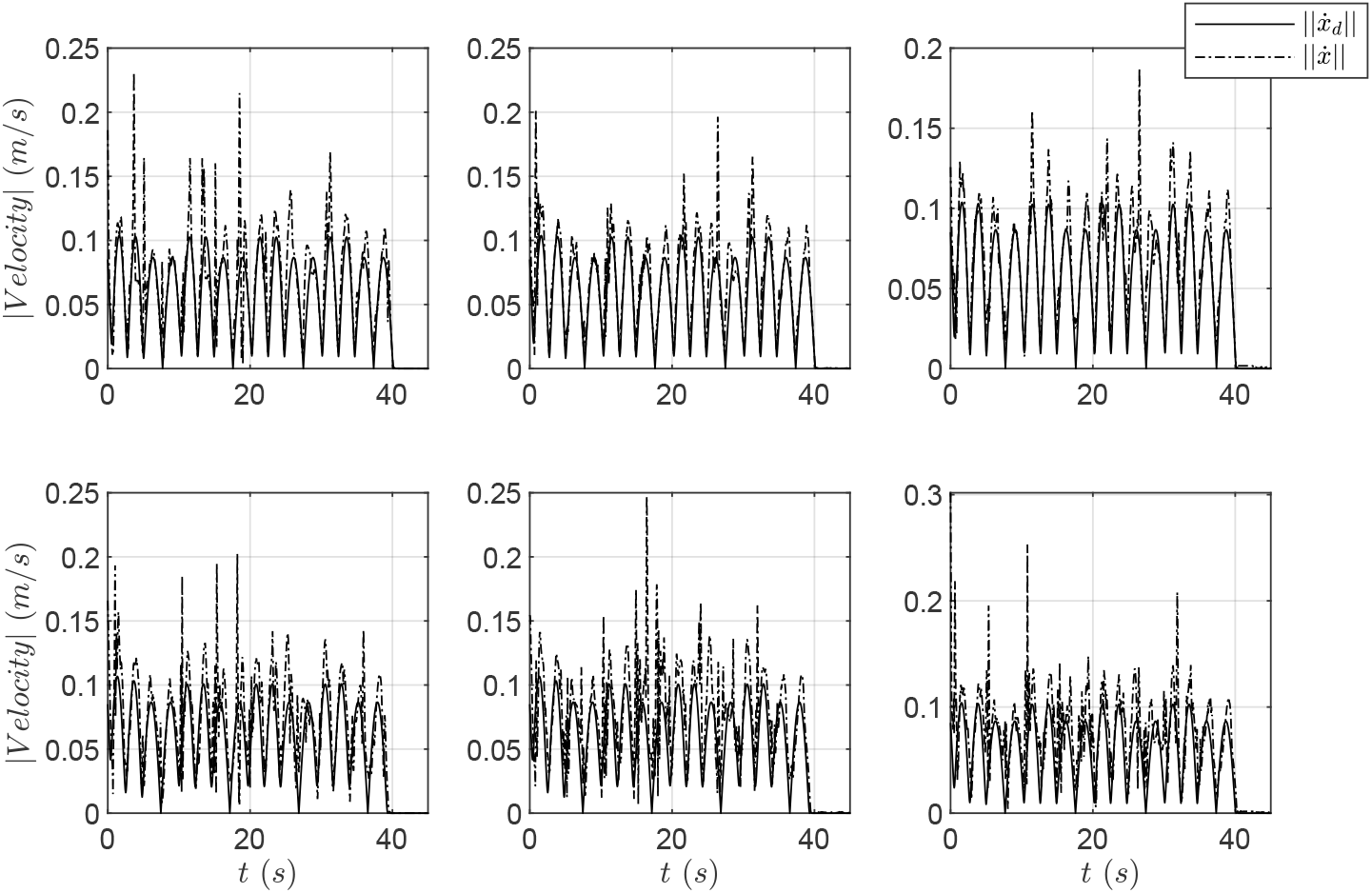
Comparing the absolute values of the desired and measured end-effector tangential velocities in the six experiments in Figure 3. The data from the experiment without delay are on the top row, the No Gravity experiment is on the left, the experiments with just gravity are on the centre, and the viscous force field results are reported on the right. The data show the characteristic bell-shaped profile; however, the measured trajectories show sporadic peaks that are probably generated by a combination of local perturbation and the limitations in motion coordination introduced by fixing the controller parameters.

## 6. Discussion

The results show that the H-PMP can generate human-like movements without the need for numerical optimisation and regression of the controller parameters from the human data. The architecture is also robust to the communication delay intrinsic to neuronal transmission. However, the results are not perfect, and they show that the proposed method has a higher variability of the *C* values compared to both human data and earlier results obtained on a 3-link arm without delays [4,40]. The data analysis suggests that this difference might be related to the need for better tuning of the architectural parameters and better coordination in issuing the motor command. Additionally, the analysis tracking errors reported in Tables I and II indicate that a state estimator is needed to reduce the accumulation of error which increases the errors for 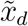 beyond the selected task’s accuracy, set with 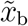.

Nevertheless, these issues mentioned above are beyond the scope of this manuscript, which focuses on the analysis of the feasibility and the stability of the proposed method. Consequently, we will continue to further investigate methods which can improve the coordination between the different architectural components and reduce systematic errors. It is worth mentioning that the data is compared against the data of young, healthy individuals [40] during the execution of a trivial reaching task. Therefore, these human subjects can be assumed to be at the best of their performances considering the health and their motor training. In fact, research data comparing hand trajectories post-stroke and healthy subjects show how the number and relative magnitude of the velocity peaks (i.e., sub-movements) are correlated with the pathological severity [49].

The performance of the H-PMP indicates that the proposed method is capable of robustly generalising movements across different environmental dynamics, and it can induce human-like movements without regression from human data. The H-PMP integrates into a single computationally inexpensive architecture the task-separation principle, the uncontrolled manifold hypothesis, and the EPH without using projected dynamics and non-linear optimisation. In contrast, the *λ*_0_-PMP and the OFC rely on numerical optimisation and projected dynamics, meaning their optimised strategies couple the system and the task dynamics, affecting the generality of the solution.

The H-PMP is composed of a series of hierarchically semi-autonomous controllers and planner and a deterministic non-linear mapping for postural optimisation robust to singularities [5, 43]. The planners and controller are stable non-linear oscillators (i.e., fractal attractor) that can be tuned online, and are robust to delays and model uncertainties [4, 5, 9, 10, 42]. Moreover, it is also an algorithmic representation of a Liénard system (i.e., Van der Pol oscillators and CPG) [5], which have been observed in biological systems [2,25,37]. The non-linear mapping used for the postural optimisation (Equation 3 and Equation 4) solves the static component of the task-separation principle, while the Task-Space Interaction Controllers (Equation 7) account for the dynamic interaction. This implies there exists a deterministic solution to the task separation principle that can be learned by neural networks generating non-linear maps. Additionally, motor strategies could be classified based on the affinity between the network input state, which agrees with the experimental observation of the motor synergies [50, 51]. These two characteristics are also compatible with the Adaptive Resonance Theory (ART), which is a neural network framework capable of explaining how the nervous system learns, classifies and adapt in changing environmental conditions by building a non-linear representation of the knowledge [52].

In regards of the dynamic primitives, they are a concept originated in robotics to describe robot control strategies regressed by the desired task-space dynamics and recently borrowed by computational neuroscience to explain motor primitives in the contest of dynamic interaction [1, 2, 14, 17, 18]. This parallel has been suggested based on the observation that humans synchronise with external dynamics [3, 50, 53], and the dynamic primitives provide a model-based method to regress the interaction with the external dynamics [1, 54]. However, this also implies that the motor primitive stability and accuracy depend on the quality of the environmental dynamics model, and they are difficult to generalise due to the coupling of the body and environmental dynamics. An equivalent form of regression of the control law from the desired task-space dynamics to solve the task-separation principle is performed by both the OFC and the *λ*_0_-PMP via numerical optimisation and projected dynamics [20, 28].

In contrast, the H-PMP deterministic mapping suggests that the dynamics primitives can be explained by exploiting the postural optimisation to achieve the desired alignment of the limb mechanical property with the principal direction of the environmental attractor. Meanwhile, trajectory desired velocity and limb stiffness modulation can be used to achieve synchronisations and desired reluctance to perturbation, respectively. This interpretation is supported by the data collected for studying how humans learn external dynamics, which shows that task performance increases with the synchronisation of the human strategy with the attractor of the external dynamics [3, 50, 55–58]. The process can be divided into three subtasks: 1) Understanding the direction of the energy flow (i.e., attractor principal directions). 2) Synchronisation with the autonomous dynamics (i.e., getting the timing right). 3) Control the tracking accuracy (i.e., impedance modulation). It should be noted that these are the same experiment, which used to justify the regression-based approach mentioned above.

## 7. Conclusion

The H-PMP explains how the nervous system can robustly control the human body in challenging dynamic conditions despite the substantial information delays. The model achieves these results without any formal modelling of either the limb and the environment dynamics. In the future, the architecture’s performance could be further improved by introducing the haptic feedback, the state estimator and learning algorithms for the online tuning and synchronisation of the architecture planner and controllers.

## 8. Acknowledgements

This work has been supported by EPSRC UK RAI Hub ORCA (EP/R026173/1), National Centre for Nuclear Robotics (EPR02572X/1) and THING project in the EU Horizon 2020 (ICT-2017-1).

## Notes

### Competing Interest Statement

The authors have declared no competing interest.

### Summary of Updates

Some minor corrections have been made to the text to clarify the manuscript. Some typos have been corrected

